# Virus-like vesicles expressing multiple antigens for immunotherapy of chronic hepatitis B

**DOI:** 10.1101/491985

**Authors:** Timur O. Yarovinsky, Stephen W. Mason, Manisha Menon, Marie M. Krady, Maria Haslip, Bhaskara R. Madina, Xianyong Ma, Safiehkhatoon Moshkani, Carolina Chiale, Anasuya Chattopadhyay Pal, Bijan Almassian, John K. Rose, Michael D. Robek, Valerian Nakaara

## Abstract

Infection with hepatitis B virus (HBV) can initiate chronic hepatitis and liver injury, eventually progressing to liver fibrosis or cancer and causing more than 600,000 deaths each year worldwide. Current treatments for chronic hepatitis B, relying on nucleoside antivirals and interferon, are inadequate and leave an unmet need for immunotherapeutic approaches. This report describes virus-like vesicles (VLV), a form of self-amplifying RNA replicons, which express multiple HBV antigens (polymerase, core, and middle surface) from a single vector (HBV-VLV). The HBV-VLV induces HBV-specific T cell responses to all three HBV antigens. Immunization of naive mice with the multiantigen HBV-VLV renders them resistant to acute challenge with HBV delivered by adeno-associated virus (AAV). Using a chronic model of HBV infection by AAV delivery of HBV, we demonstrate immunotherapeutic potential of the multiantigen HBV-VLV in combination with DNA booster immunization, as 40% of the HBV-VLV-treated mice showed a decline of the serum HBV surface antigen below the detection limit and marked reduction in liver HBV RNA accompanied by induction of HBsAg-specific CD8 T cells. These results warrant further evaluation of multiantigen HBV-VLV for immunotherapy of chronic hepatitis B.

**IMPORTANCE:** More than 240 million people worldwide are chronically infected with hepatitis B virus. Current therapies are not sufficiently effective and are often beyond reach in the developing world. We describe a virus-like vesicle-based immunotherapeutic vaccine that expresses three major antigens of hepatitis B virus as a self-amplifying RNA replicon. By incorporating three HBV antigens in a single vaccine, we ensure broad T cell responses. We demonstrate that immunization with this vaccine protects mice from hepatitis B virus in a model of acute challenge. Importantly, treatment with this vaccine shows 40% efficacy in a mouse model of chronic hepatitis B. Thus, this study paves the way for evaluation of the multi-antigen virus-like vesicles as a tool for immunotherapy of chronic hepatitis B.

## Introduction

Human hepatitis B virus (HBV), a partially double stranded DNA virus of the genus Orthohepadnavirus, replicates in hepatocytes and causes liver inflammation and injury (1–3). While most adults recover from the acute infection and become immune to subsequent exposures, 2-6% of adults and 90% of infants develop chronic hepatitis B (CHB) and are at increased risk of developing liver cirrhosis or liver cancer (2, 3). Multiple factors, including vertical transmission to newborns before the immune system is fully formed and active suppression of the immune system by hepatitis B surface antigen (HBsAg) and hepatitis B e antigen (HBeAg), contribute to HBV persistence (4). Worldwide, 240-350 million people are chronically infected with HBV and 600,000-900,000 die each year from HBV-associated liver diseases (5). In the United States, CHB prevalence is estimated at 850,000 and the rate of new infections remains about 1 per 100,000 population despite routine use of the prophylactic HBV vaccine (6). Currently approved therapeutic approaches for CHB, including pegylated interferon-α (IFNα) and nucleos(t)ide analogues, may reduce and control HBV replication in a subset of patients (3). However, adverse effects of IFNα treatment, lack of complete HBV clearance, and development of drug resistance after prolonged treatment with nucleos(t)ides emphasize the unmet need for alternative therapeutic approaches that can lead to functional cure (3, 7, 8).

Spontaneous clearance of HBV through immune-mediated mechanisms, particularly T cell mediated immune responses, in acute and in some chronically infected patients provides a rationale for CHB immunotherapy (9). Ongoing efforts at preclinical and early clinical stages include activation of innate immunity through TLR ligands, treatment with immune checkpoint inhibitors, and/or immunization with HBV antigens delivered by plasmid DNA or viral vectors (7, 9). DNA immunization is a safe and reliable method for induction of antibody responses, but this approach requires special delivery such as electroporation, and remains poorly immunogenic for T cells (10). On the other hand, viral vectors are highly immunogenic and efficient at mounting T cell responses, but their safety remains a concern even for attenuated and replication-deficient viral vectors (9).

Virus-like vesicles (VLVs), a form of self-amplifying RNA replicons, are a non-pathogenic and immunogenic alternative to viral vectors. VLVs have an excellent safety profile and induce T cell and antibody responses (11–13). The RNA-dependent RNA polymerase from Semliki Forest virus and glycoprotein from vesicular stomatitis virus (VSV-G) expressed from the same vector are essential components of the VLV platform that enables rapid and efficient production of VLVs in BHK-21 cells and expression of the desired antigens in the immunized host (13). A previous study showed that immunization with VLVs expressing HBV middle surface antigen (MHBs) induced HBV-specific CD8^+^ T cells and protected mice against experimental acute HBV infection (14). Furthermore, HBV-transgenic mice with low HBeAg produced HBV-specific CD8^+^ T cells after priming with plasmid DNA and boosting with the VLV expressing MHBs, suggesting that VLV immunization may break immune tolerance, a characteristic feature of CHB (14). While that study demonstrated promise for VLV-based immunotherapy, it is evident from other studies that immunization with multiple HBV antigens to enhance and/or broaden T cell responses is necessary for therapeutic effect and HBV elimination (15–17).

In this report, we describe the design, production, immunogenicity, and immunotherapeutic potential of polycistronic VLV vectors that express three HBV antigens: polymerase (Pol), core (HBcAg), and MHBs. Our goal was to target multiple HBV antigens simultaneously and to elicit the broad and vigorous multi-specific immune responses needed for the resolution of CHB infection in humans. To that end, we co-expressed structural (MHBs and HBcAg) and non-structural (Pol) HBV proteins as a single polycistronic open reading frame. This approach enabled the induction of broad T cell responses to multiple antigens that were sufficient to protect mice from experimental acute HBV replication. In a murine model of experimental CHB driven by AAV-HBV transduction, prime immunization with the multiantigen VLV vector followed by two booster DNA immunizations induced HBsAg-specific CD8^+^ T cell responses and reduced serum HBsAg and liver HBV RNA.

## Results

### VLV design and production

Since HBV Pol and HBcAg are important targets for CD8^+^ T cells (18–20), we aimed to incorporate these antigens into the new VLV vectors for immunization. The previously reported VLV vector (referred to as MT2A and depicted in Fig 1A) relied on the ribosome-skipping 2A peptide from *Thosea asigna virus* (T2A) to express both MHBs and VSV-G from the same subgenomic RNA (14). Using that vector as a backbone, we designed and constructed VLVs for expression of HBV polymerase alone (PolT2A) or as a polycistronic vector for expression of Pol, HBcAg, and MHBs by interspacing the HBV antigens with the ribosome skipping peptides (3xT2A) as depicted in Fig 1A. We replaced several key amino acids in the terminal protein domain of Pol (Y63A, W74A, Y147A and Y173A) to abolish its DNA binding (primase) activity (21). In addition, we deleted amino acids 538-544 encompassing the active site (residues YMDD) of the reverse transcriptase domain to remove its RNA-dependent DNA polymerase and DNA-dependent DNA polymerase activities. Furthermore, we introduced mutations into the RNAse H domain (amino acids 680-832) in order to abolish the ribonuclease H activity, as previously described (22). We altered the HBcAg sequence to improve processing and peptide presentation by substituting the disulfide bond-forming cysteine residues (C48S, C61S, C107S and C183S) and adding an N-terminal fusion with the 76-amino acid ubiquitin (Ub) peptide (20, 23). To produce VLVs for evaluation of antigen expression and immunogenicity, we transfected BHK-21 cells with the plasmid DNA expressing 3xT2A VLV RNA under the control of CMV promoter, collected the master VLV stocks and used them for propagation and concentration of VLVs by ultrafiltration. The resulting VLV titers routinely exceeded 10^9^ pfu/ml.

**Figure 1.**
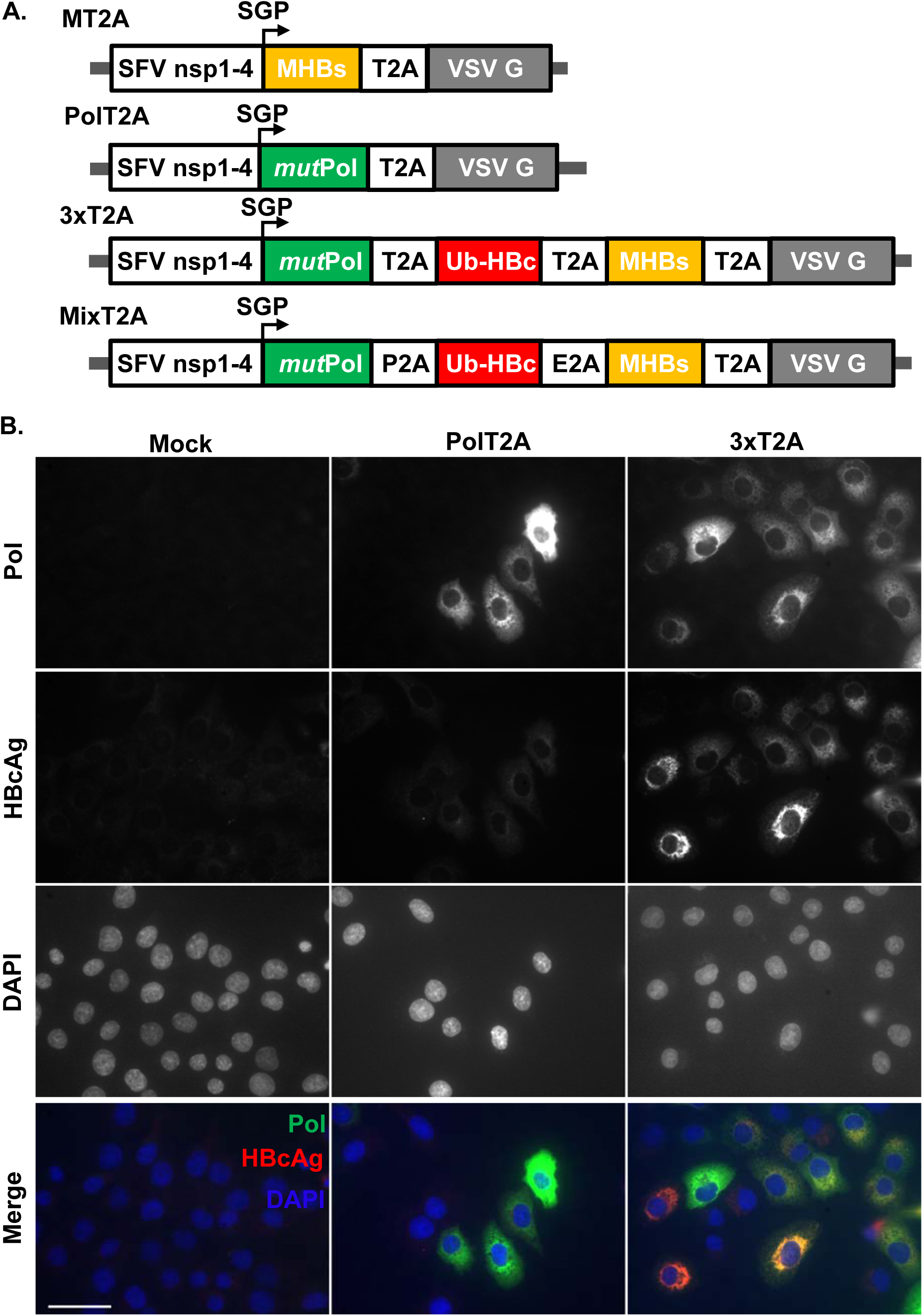
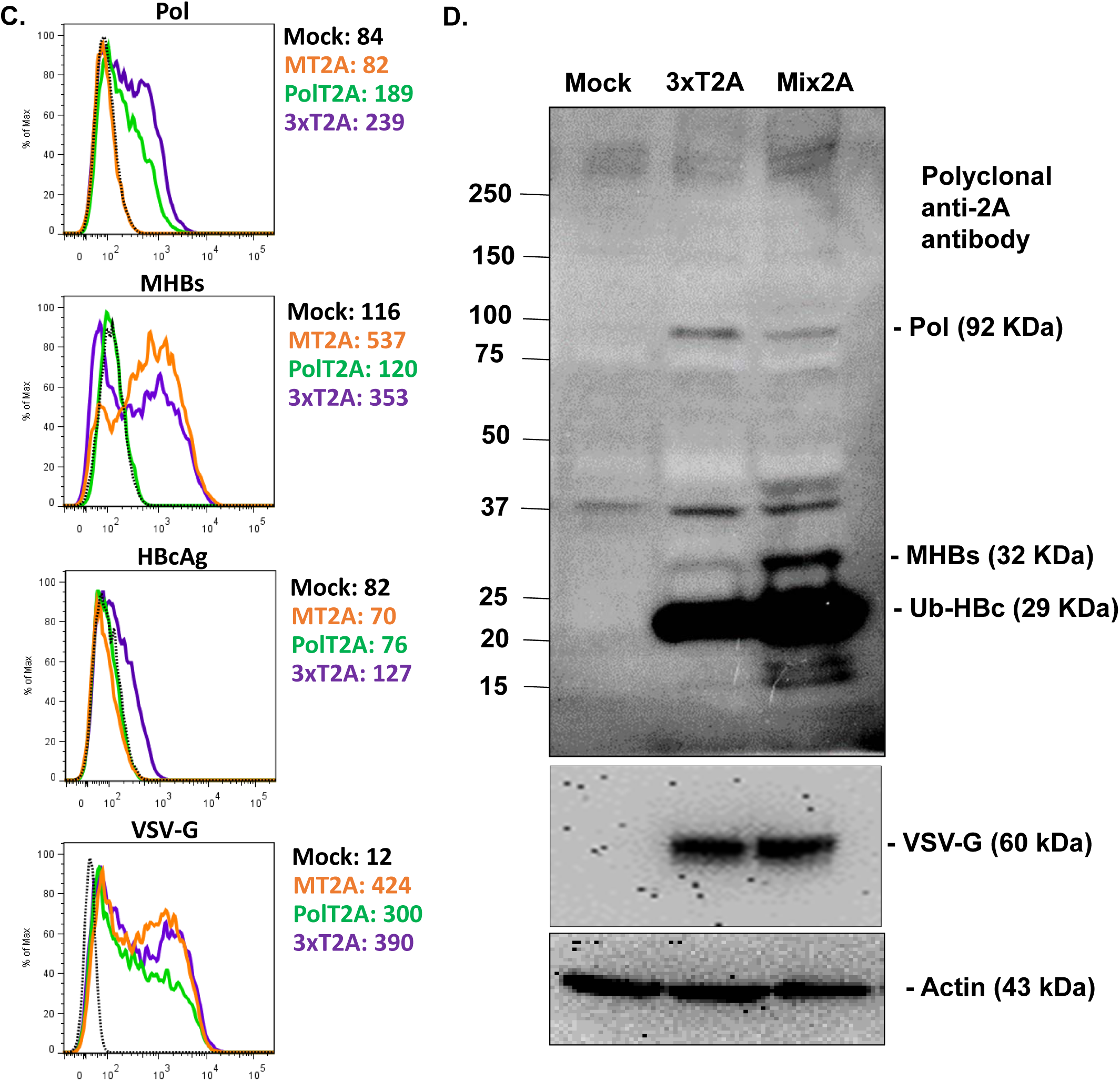
Virus-like vesicle platform for therapeutic HBV vaccine expressing polymerase (Pol), ubiquitinated core (Ub-HBc), and middle surface (MHBs) antigens. (A) Design of the replicating VLV for expression of MHBs (MT2A), HBV Pol (PolT2A) single antigens and for polycistronic expression of the three HBV antigens using different 2A self-cleaving peptides (3xT2A and Mix2A). (B) Validation of antigen expression in BHK-21 cells after infection with PolT2A or 3xT2A VLV (MOI=1) by immunofluorescence at 20 h post infection using the antibodies specific for HBV Pol and HBcAg. (C) Validation of antigen expression in BHK-21 cells after infection the VLV expressing MHBs only (MT2A), Pol only (PolT2A) or the three antigens (3xT2A) by flow cytometry using the antibodies specific for Pol, MHBs, HBcAg, and VSV-G. Geometric mean fluorescence intensity is shown for each antibody and VLV. (D) Evaluation of antigen expression and VLV replication in BHK-21 cells transfected with plasmid DNA for 3xT2A or Mix2A VLV using the 2A-peptide specific antibody in western blots. Anti-VSV-G and anti-actin were also used as controls. The data are representative of three independent experiments.

To mitigate potential risks of homologous recombination of the repeating T2A peptide sequences in the 3xT2A construct during VLV replication, we designed and evaluated an alternative construct, Mix2A, in which we replaced the two T2A peptides with the porcine teschovirus-1 (P2A) and equine rhinitis A virus (E2A) peptides (24) (Fig 1A). These peptides are structurally similar, but encoded by different nucleotide sequences. When produced in BHK-21 cells under identical conditions, the resulting titers were similar for Mix2A VLVs (2.2*109 pfu/ml) and 3xT2A VLVs (1.8*109 pfu/ml).

### Validation of antigen expression

We evaluated expression of Pol and HBcAg in BHK-21 cells infected with PolT2A and 3xT2A VLVs by immunofluorescence (Fig 1B). Using HBV Pol specific mAb clone 2C8 (25), we observed a characteristic fine granular staining in the cytoplasm at 24 h after infection with either of the VLVs (Fig 1B, top row). Staining with the HBcAg-specific polyclonal antibody showed variable levels of HBcAg expression in most of the BHK-21 cells infected with 3xT2A, but not with the PolT2A VLV (Fig 1B, second row).

To compare expression levels of HBV antigens in BHK-21 cells infected with polycistronic VLVs 3xT2A versus VLVs expressing Pol or MHBs alone, all at MOI=1 at 24 h post infection, we used intracellular staining with the antibodies for Pol, HBcAg, MHBs, followed by flow cytometry analyses (Fig 1C). The cells infected with 3xT2A VLVs showed slightly higher levels of Pol expression, compared to PolT2A VLVs. On the contrary, cells infected with 3xT2A VLVs showed slightly lower levels of MHBs expression compared to MT2A VLVs. As expected, no Pol staining was detected in cells infected with the MT2A VLVs and no MHBs staining was detected in cells infected with PolT2A VLVs. Staining for HBcAg was detectable only in cells infected with 3xT2A, albeit at relatively low levels. It is important to note that VSV-G, the only structural component of VLVs, was detectable in most BHK-21 cells after infection with all three VLV vectors above (Fig 1C).

We also compared antigen expression from 3xT2A or Mix2A VLVs by immunoblotting of BHK-21 cell lysates at 24 h post-infection at MOI=1 (Fig 1D). Using a polyclonal anti-2A peptide antibody, which recognizes the GDVESNPG epitope shared by all 2A peptides, we detected bands corresponding to the predicted molecular weights of HBV polymerase (92 kDa), middle S antigen (32 kDa) and modified core (29 kDa), which is consistent with the expectation that each of the antigens is tagged with the 2A-peptide sequences at the C-terminus. We also observed additional bands at ∼16, 18, and 40 kDa in the lysates from Mix2A VLV-infected cells, which may be due to incomplete recognition of the 2A peptide sequences during translation or subsequent proteolytic degradation. In addition, we noted the high molecular weight (>250 kDa) and a 37-kDa non-specific bands present in lysates from non-infected and VLV-infected cells, which most likely are the result of the antibody cross reactivity with the endogenous proteins in BHK-21 cells. VSV-G was detected by polyclonal anti-VSV-G antibody only in the VLV-infected cells with comparable intensity between 3xT2A and Mix2A.

### Immunogenicity of 3xT2A and Mix2A VLV vaccine candidates in naïve mice

Administration of 3xT2A or Mix2A VLVs at 10^8^ pfu/mouse resulted in an apparent increase in frequency (Fig 2A) and absolute numbers (Fig 2B) of CD44^hi^CD62L^lo^ effector CD4^+^ and CD8^+^ T cells in spleens at day 7 post-immunization. Furthermore, we quantified the frequencies (Fig 2C) and absolute numbers (Fig 2D) of CD8^+^ T cells that produce IFNγ after ex vivo restimulation with the peptides derived from the epitopes from within HBV Pol, HBcAg, and MHBs. The cumulative frequencies of the IFNγ^+^ (i.e., HBV-specific) CD8^+^ T cells exceeded 2%; among those the CD8^+^ T cells that recognized the HBS-371 peptide were most prevalent. Furthermore, substantial numbers of CD8^+^ T cells responded to HBV peptide stimulation by producing IFNγ and TNF (Fig 2E and 2F), indicating that VLV may induce double-cytokine producing CD8^+^ T cells, which are efficient in non-cytolytic elimination of HBV from the infected hepatocytes (26). The 3xT2A vector showed a trend towards higher frequency and absolute numbers of MHBs-specific CD8^+^ T cells when compared to Mix2A, albeit with no statistically significant difference. Since the use of different 2A peptides in the Mix2A VLV construct did not provide any apparent advantage for HBV antigen immunogenicity in naïve mice, we proceeded to use the simpler of the two constructs, 3xT2A, for efficacy studies.

**Figure 2.**
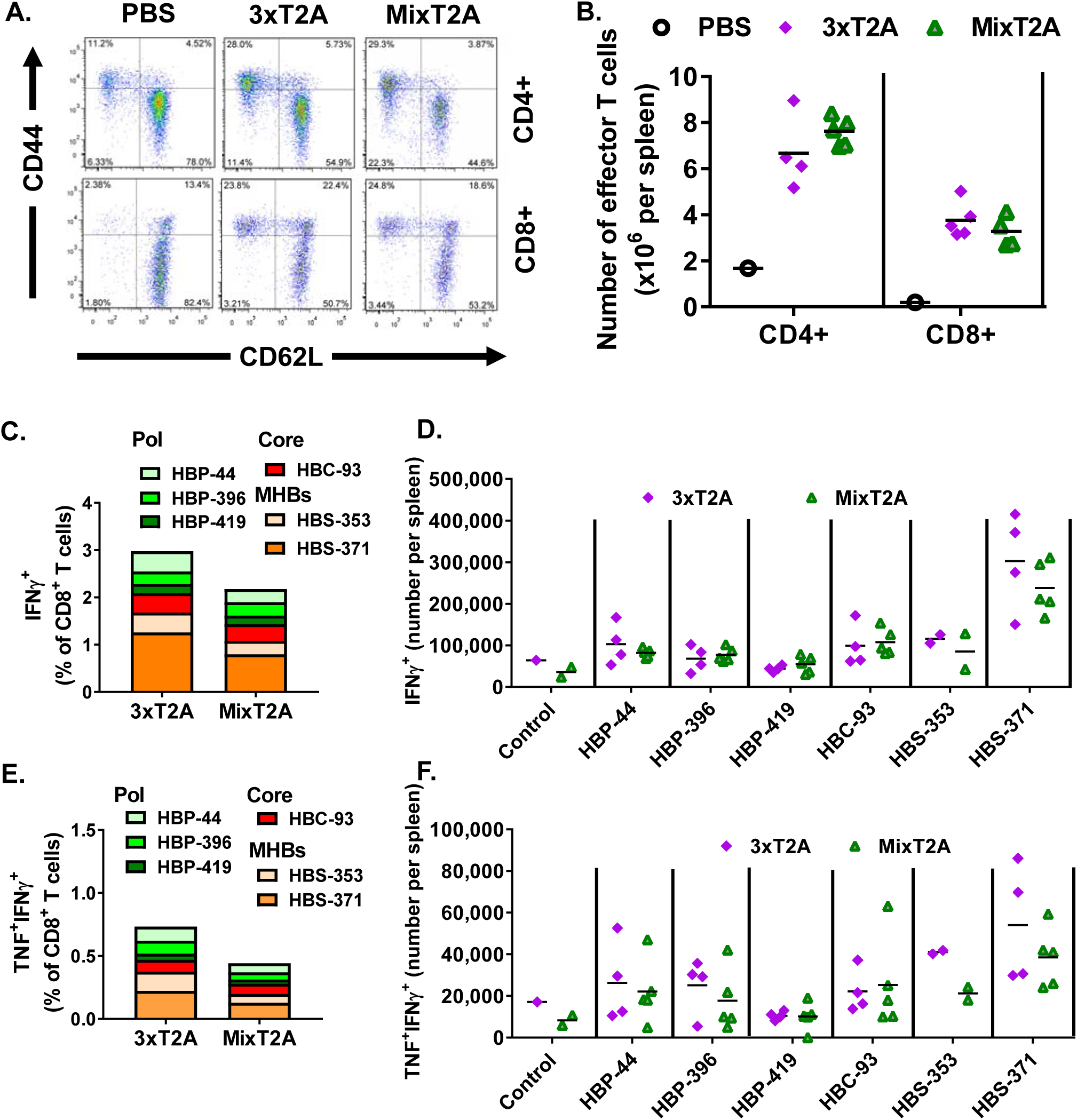
Comparison of immunogenicity of the 3xT2A and Mix2A VLV in naïve C57BL/6J mice. Accumulation of effector CD4^+^ and CD8^+^ T cells at day 7 after immunization with 3xT2A or Mix2A VLV. (A) Representative plots for CD62L and CD44 staining after gating on CD4^+^ or CD8^+^ T cells. (B) Absolute numbers of effector (CD44hiCD62Llo) T cells. (C) Frequency of HBV-specific IFNγ-producing CD8^+^ T cells identified after ex vivo stimulation with the indicated peptides from Pol (HBP-44, HBP-396, HBP-419), HBcAg (HBC-93) and surface (HBS-353 and HBS-371) quantified by intracellular staining and flow cytometry. (D) Numbers of IFNγ^+^ CD8^+^ T cells per spleen. (E) and (F) Frequency and numbers of HBV-specific double cytokine (TNF+IFNγ^+^)-producing CD8^+^ T cells, respectively. Values in C and E are mean of 3 to 5 mice per group.

### Immunization with 3xT2A VLV protects mice from experimental acute challenge with AAV-HBV

To determine whether VLV-mediated expression of HBV Pol, HBcAg and MHBs generates protective immunity against HBV, we immunized CB6F1/J mice with 3xT2A or empty vector VLV and administered an equivalent volume of PBS to a control group of mice. Six weeks later, we challenged these mice with HBV delivered to the liver via recombinant adeno-associated virus (AAV-HBV) encoding a 1.3-overlength HBV genome and monitored serum HBsAg and HBeAg as a measure of HBV replication for additional 4 weeks. HBsAg levels peaked in PBS and empty vector VLV control groups of mice 14 days after infection, but remained low in 3xT2A VLV-immunized mice throughout the experiment (Fig 3A). HBeAg levels steadily increased during infection in the PBS and empty vector VLV groups (Fig 3B), but barely rose above background at days 14 and 21 and declined by day 28 in 3xT2A VLV-immunized mice. Quantitative analyses of HBV RNA in the livers of mice euthanized on day 28 showed statistically significant decrease in the 3xT2A VLV immunized group compared to empty vector VLV group (Fig 3C). 3xT2A VLV-immunized mice, but not the mice immunized with the empty vector VLV, showed a statistically significant increase in HBsAg-specific cells in the spleens that produce IFNγ after stimulation with the dominant peptide epitopes (Fig 3D). Thus, immunization with the 3xT2A VLV expressing the HBV Pol, HBcAg and MHBs protects mice from experimental acute HBV replication.

**Figure 3.**
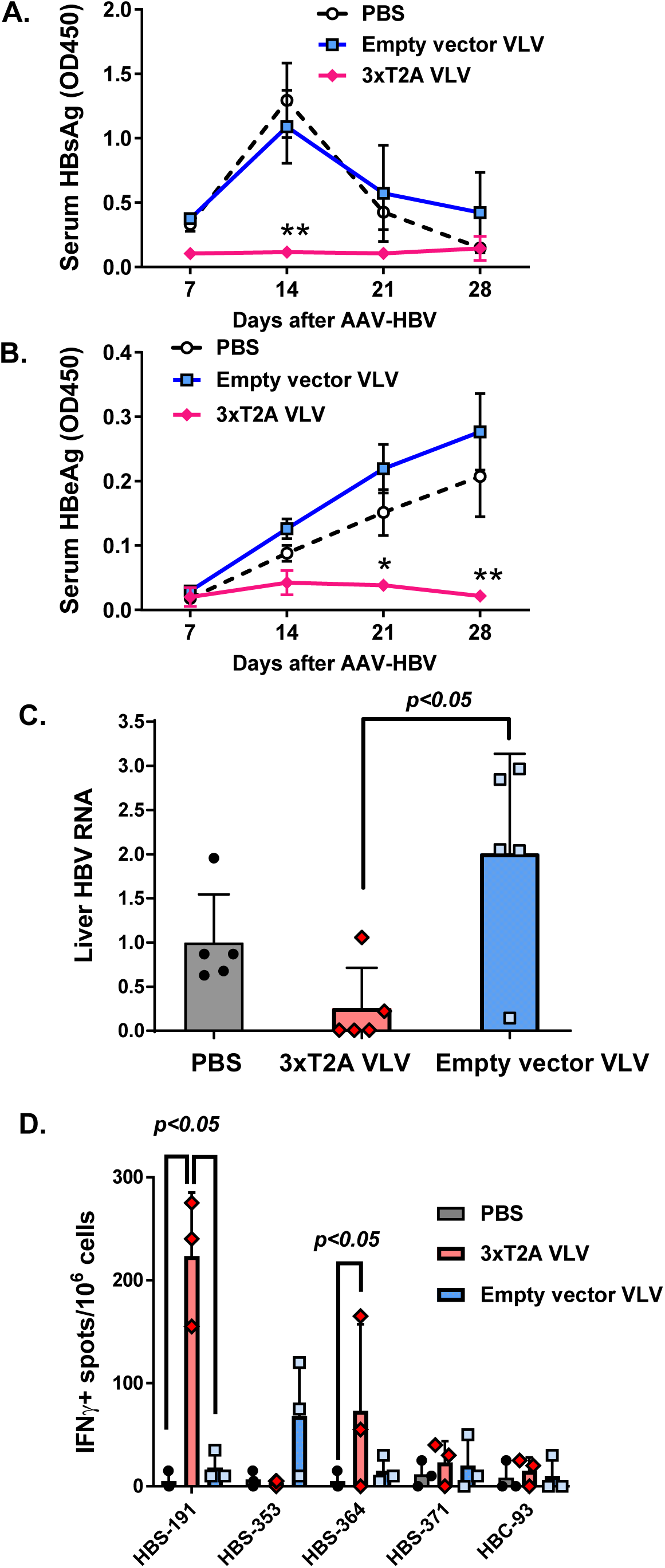
Efficacy of 3xT2A VLV as prophylactic vaccine in acute HBV infection. CB6F1 mice were immunized with 3xT2A VLV or empty vector VLV for 6 weeks prior to challenge with AAV-HBV. (A) Levels of serum HBsAg (OD450) measured by specific ELISA from blood samples taken weekly. (B) Weekly levels of HBeAg. The values are mean±SEM (n=5). Asterisks indicate significant difference (*, *p*<0.05, **, *p*<0.01) in 3xT2A-immunized mice compared to PBS or empty vector VLV. (C) Abundance of HBV RNA in liver samples from mice euthanized on day 28 after challenge with AAV-HBV. TaqMan-based qRT-PCR was used to quantify HBV RNA and mouse GAPDH mRNA. The HBV RNA data were normalized to GAPDH and expressed as relative to the mean in the PBS group. Individual values and mean±SD are shown. (D) Frequency of HBV-specific IFNγ^+^ producing cells was measured by ELISPOT on day 28 after challenge with AAV-HBV after ex vivo stimulation of splenocytes with the indicated peptides. Individual values and mean+SD are shown.

### Evaluation of 3xT2A VLV in an experimental model of chronic HBV infection

We explored the therapeutic potential of 3xT2A in the setting of the AAV-HBV mouse model of chronic HBV infection (27). To initiate this model, we transduced C57BL/6J male mice with AAV-HBV-1.2 (2*10^11^ genome copies of AAV-HBV particles per mouse) and monitored serum HBsAg levels every week. As expected, we observed a steady rise of serum HBsAg from 11.52±2.40 ng/ml (mean±SEM, n=20) at day 7 to 70.04±6.25 ng/ml (mean±SEM, n=20) at day 21 post AAV-HBV transduction. Based on the measurements at day 21, we formed 2 groups of 10 mice each with equivalent levels of serum HBsAg prior to immunization with 3xT2A or empty vector VLV on day 28 post AAV-HBV transduction. Fig 4A shows responses to the treatment through the full course of the experiment as the fold change of HBsAg levels (OD450) relative to day 28, whereas Fig 4B shows the concentrations of serum HBsAg in ng/ml after VLV prime. Both VLV vectors caused an apparent, but transient, decline in HBsAg levels at day 35, which may reflect host interferon response to VLVs. HBsAg levels rebounded in both groups by day 42 and remained essentially steady for the empty vector VLV group. By day 49, the average HBsAg levels of the 3xT2A VLV-immunized groups started to decline and become significantly lower at days 56 and 63. Importantly, by day 63, the concentration of HBsAg dropped below the limit of detection (3.12 ng/ml) in four out of ten 3xT2A VLV-immunized mice. These results indicate that 40% of the chronically infected animals responded to the treatment with 3xT2A VLV.

**Figure 4.**
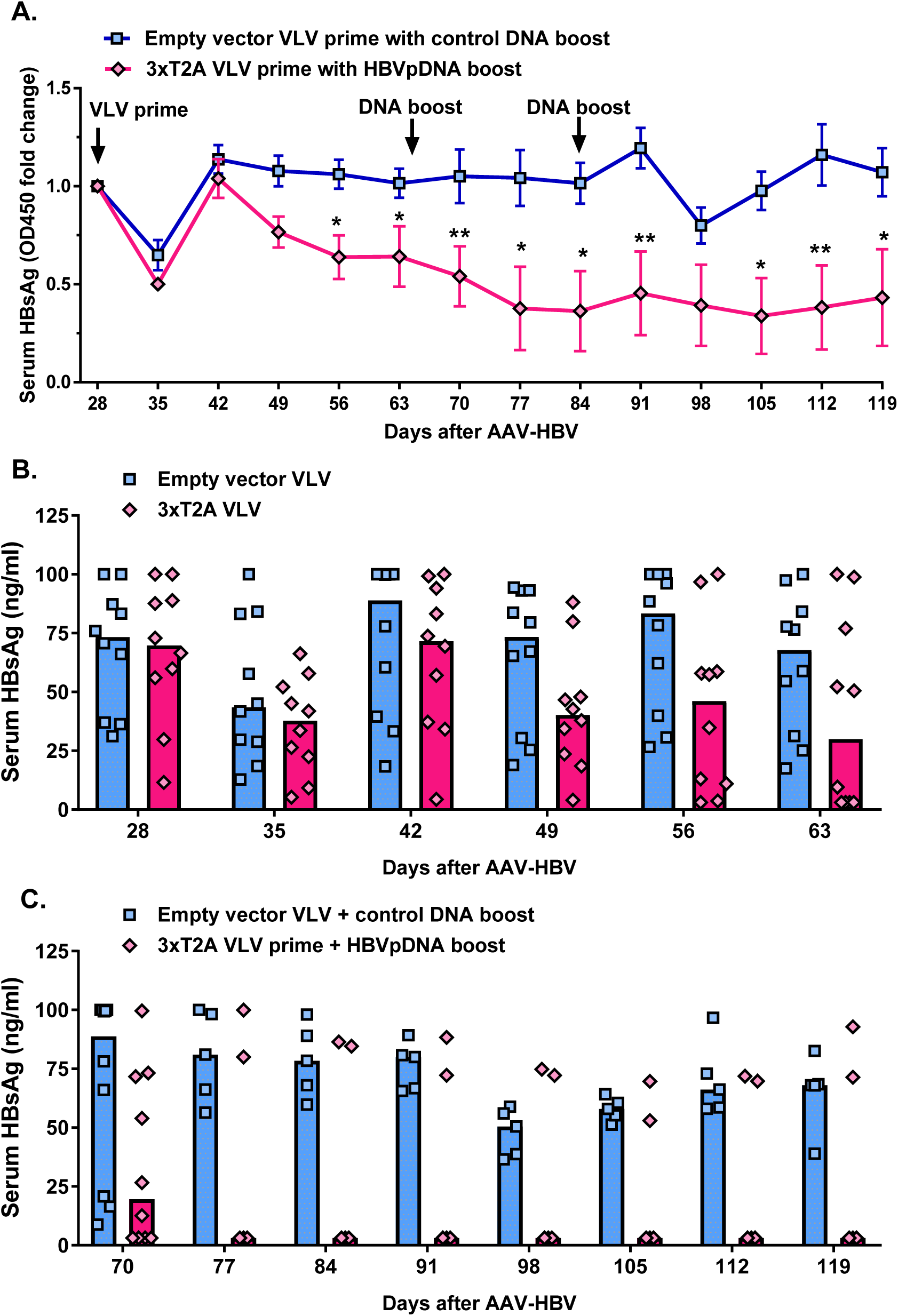
Effects of 3xT2A VLV on HBsAg levels in chronic AAV-HBV model. Persistent HBV replication was established in C57BL/6 mice by delivery of HBV genome with AAV. The groups were balanced for HBsAg prior to immunization with empty vector VLV or 3xT2A at day 28 after AAV-HBV transduction. Booster DNA immunizations were carried out on days 65 and 84, as indicated by arrows, using a pool of 3 plasmids expressing MHBs, HBcAg, and HBV Pol for 3xT2A group or empty plasmid vectors for the empty vector VLV. (A) Fold change in HBsAg levels (OD450). Values are mean±SEM (n=10 for days 28-70 and n=5 thereafter). Quantification of HBsAg (ng/ml) after VLV prime at day 28 (B) and DNA boost at day 65 (C). Individual values and median are shown by symbols and bars, respectively. Asterisks in A indicate significant difference between 3xT2A and empty vector VLV-immunized animals (*, *p*<0.05, **, *p*<0.01).

Since only 40% of the animals showed responses to 3xT2A VLV and considering that a single VLV immunization would be unlikely to overcome HBV tolerance in CHB, we added DNA immunization as a boost after blood collection on days 65 and 84 (Fig 4A). We administered a pool of three DNA plasmids expressing MHBs, HBcAg, and HBV Pol (further referred as HBVpDNA) to the group of mice immunized with 3xT2A and empty vector plasmid to the group of mice immunized with empty vector VLV. HBsAg levels continued to decline in the group of mice treated with 3xT2A VLV and HBVpDNA and, with exception of a single time point (day 98), were significantly lower (p<0.05) than in the group of mice treated with empty vector VLV and DNA (Fig 4C).

To determine whether 3xT2A VLV-induced decreases in serum HBsAg were accompanied by reduction of HBV RNA in liver, we euthanized 5 animals in each group to collect liver samples on day 70 and the remaining 5 animals at the termination of the experiment on day 119. Quantitative analyses of HBV RNA showed no difference between the groups at day 70 (Fig 5A). However, at day 119 3 out of 5 mice primed with 3xT2A VLV and boosted twice with HBVpDNA showed more than 100-fold reduction of liver HBV RNA. Remarkably, this group showed a very strong correlation between serum HBsAg and liver HBV RNA (R^2^=0.905, p=0.013 for 3xT2A VLV plus HBVpDNA vs. R^2^=0.179, p=0.478 for empty vector VLV plus control DNA), suggesting that the observed decrease of serum HBsAg is due to a reduction of HBV replication.

**Figure 5.**
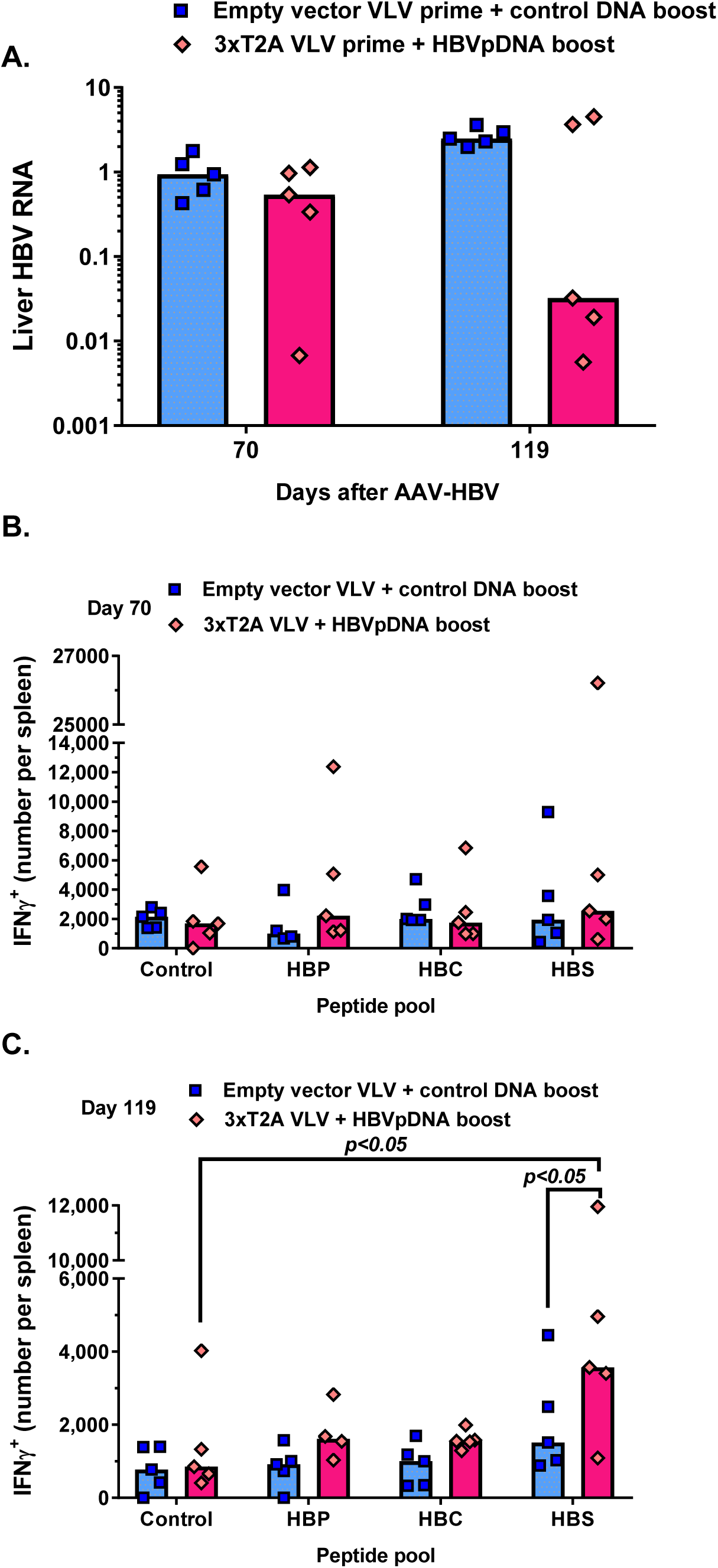
Effects of 3xT2A VLV on liver HBV RNA in chronic AAV-HBV model. Liver samples were collected at day 70 or day 119 from the animals immunized as shown in Fig 4 for qRT-PCR analyses. (A) Abundance of HBV RNA was determined after normalization to mouse GAPDH mRNA and expressed relative to the mean of the empty vector VLV group (n=5). Numbers of IFNγ^+^ CD8^+^ T cells per spleen at day 70 (B) and day 119 (C) were determined by intracellular flow cytometry after stimulation with HBV antigen peptides.

Finally, we analyzed HBV-specific CD8^+^ T cell responses in the immunized groups. At day 70, stimulation of splenocytes with HBV Pol and HBcAg peptides did not lead to an increase of IFNγ-producing CD8^+^ T cells above background (no peptide control) with the exception of a single mouse immunized with 3xT2A VLV (Fig 5B). Stimulation with HBsAg peptides revealed that the same mouse and another mouse immunized with the empty vector VLV group had IFNγ-producing CD8^+^ T cells above background. At day 119, two out of five mice immunized with 3xT2A VLV had HBsAg-specific IFNγ-producing CD8^+^ T cells in numbers that were above background, with overall statistically significant increase over the no peptide control and the empty vector control VLV (Fig 5C).

## Discussion

Our results demonstrate the feasibility and potential of the VLV platform for polycistronic expression of HBV antigens and immunotherapy for CHB. The HBV antigens expressed by 3xT2A and Mix2A VLV constitute three of the four major proteins expressed by HBV: the entire polymerase protein (HBV pol) with mutations to inactivate reverse transcriptase activity, ubiquitin-tagged core (UbHBc) with mutations to prevent the formation of stable disulfide-mediated dimers, and the middle surface antigen (MHBs). The combination of three different 2A peptides in Mix2A VLVs did not provide any obvious advantage over repeating T2A peptides in 3xT2A VLVs. The selected 3xT2A VLVs replicate to high titers (>10^9^ pfu/mL), express the individual HBV proteins of interest (Fig 1), and induce slightly higher specific CD8^+^ T cell responses to each antigen than Mix2A VLV (Fig 2). Taking advantage of mouse strain-specific differences in persistence of AAV-mediated transduction of HBV (28), we demonstrated that a single immunization with 3xT2A VLVs provides protection of CB6F1 mice from an acute challenge (Fig 3). Using the AAV-HBV-transduced mice as a model of persistent HBV replication, we evaluated the utility of the 3xT2A VLV for CHB immunotherapy. Since persistent HBV replication in liver leads to tolerance to HBV antigens (27), we did not expect that a single immunization with 3xT2A VLV would be sufficient to clear the virus. Thus, we used 3xT2A VLV for prime immunization followed by 2 booster immunizations with DNA plasmids expressing the same antigens. This immunization schedule resulted in sustained decline of serum HBsAg and liver HBV RNA in a subset of mice with established chronic HBV replication (Fig 4 and Fig 5A). These data indicate that 3xT2A VLVs function not only as a prophylactic vaccine but could also be used as a component of comprehensive immunotherapy.

Immunotherapeutic approaches for CHB, including DNA vaccination and therapeutic vaccines based on viral vectors, promise to provide functional cure by reducing or eliminating the virus from infected hepatocytes (7, 9). However, none of the experimental immunotherapies tested to date have shown sufficient efficacy in clinical trials (9). This suggests that these vaccines either lacked sufficient potency, were not targeting the correct antigen(s), were targeting too few antigens or, perhaps, all the above. It appears that success depends on the induction of CD8^+^ T cells that are both cytolytic and non-cytolytic in action rather than induction of humoral responses, as does the present prophylactic vaccines (9, 29). While therapeutic vaccines using recombinant VSV or modified vaccinia virus have a potential to induce more robust T cell mediated immune responses to HBV (30), the concerns about their safety profile may represent an obstacle for clinical applications (9).

Immunotherapeutic vaccines expressing HBV Pol, HBcAg and MHBs using the VLV vectors address the safety concerns and provide sufficient efficacy by expressing multiple HBV antigens that are optimized for induction of T cell responses. Restriction of VLV replication by type I interferons enables 100% survival rate of mice even after intracranial administration of VLV (31). Inclusion of HBV Pol provides a number of highly immunogenic and conserved epitopes for CTL that can detect and eliminate HBV-infected cells and limit viral spread (18). Since expression of intact (wild-type) HBcAg from a single antigen-expressing VLV did not induce sufficient T cell responses to control HBV replication (14), the polycistronic constructs described in this report included ubiquitinated HBcAg with mutated cysteine residues. These changes may have improved peptide presentation to T cells through prevention of HBcAg dimerization and enhancement of proteolytic degradation (20). Even though MHBs is expressed in the third position in the 3xT2A and Mix2A vectors, it appeared as the dominant antigen, as demonstrated by the frequency of IFNγ^+^ and IFNγ^+^TNF^+^ CD8^+^ T cells after immunization (Fig 2C and Fig 2D). Thus, the polycistronic design of the VLV vector induces broad T cell responses in naïve mice with a potential to eliminate HBV through cytolytic and non-cytolytic mechanisms.

To evaluate prophylactic and therapeutic efficacy of 3xT2A VLV, we employed both acute and chronic models of HBV infection established by delivery of HBV genome DNA via liver-tropic AAV (27). This approach overcomes the species-specific restrictions in early steps in the HBV replication cycle and enables homogenous transduction of mouse hepatocytes, HBV replication, and secretion of HBsAg (27). Delivery of HBV via AAV also provides more reproducible results than hydrodynamic delivery of HBV genome in plasmid DNA for acute challenge. While the HBV-specific T cell responses are muted as the chronicity of infection causes T cell exhaustion and tolerance to HBV antigens, the mechanism and level of tolerance are different from those in the HBV transgenic mouse model (27, 28, 32). Thus, despite some limitations, including highly variable levels of viral load and HBsAg levels, the AAV-HBV model is more suitable for evaluation of immune-mediated clearance of HBV.

After determining that 3xT2A VLV can protect mice from acute AAV-HBV challenge (Fig 3), we evaluated the immunotherapeutic potency of 3xT2A in a mouse model of CHB. As a precaution that a single immunization with 3xT2A VLV might not be sufficient to break tolerance in the chronically infected mice, we added DNA booster immunization to the treatment protocol. We observed that priming of chronically infected mice with 3xT2A VLV, followed by booster DNA immunization, results in a statistically significant decline of serum HBsAg (Fig 4) with 40% of the mice dropping the HBsAg levels to the limit of detection and showing marked reduction of liver HBV RNA (Fig 5A). Such a partial response is not unheard of with immunotherapeutic approaches, and may be due to variable levels of HBV replication or differences in the TCR repertoire of T cells responding to the viral antigens. Following this immunization schedule, a subset of mice also showed a statistically significant accumulation of HBsAg-specific IFNγ-producing CD8^+^ T cells in the spleens at later time points (Fig 5C). Although the design of the study described here did not allow us to be certain that the booster DNA immunizations would be necessary, our data clearly indicate that the multiantigen HBV-VLV shows promise as an immunotherapeutic tool for treatment of CHB.

Among other therapeutic vaccines for CHB evaluated in the HBV-AAV model, the adenovirus serotype 5 (Ad5)-based therapeutic vaccine (TG1050) is comparable to our VLV-based vector in approach and goals of inducing HBV-specific T cell responses (16). TG1050 expresses a fusion protein consisting of a deleted and mutated HBVpol, truncated HBcAg, and two regions of HBsAg (16). Similar to our observations with 3xT2A VLVs, the TG1050-treated group of mice showed a three-fold reduction in HBsAg levels compared to those in the control group. While the effects of TG1050 on individual animals were not reported, we observed that 40% of the animals in our study responded by decline of HBsAg to the limit of detection. TG1050 presently is in Phase I/II clinical trials (NCT02428400) in HBV-infected patients with the results of this trial expected in 2019.

In order to improve immunogenicity and efficacy of therapeutic vaccine candidates for chronic HBV infection, others have used a combination of approaches, such as adjuvants, addition of checkpoint inhibitors, and prime/boost regimens (7, 9). Recent studies showed that an optimized, heterologous prime/boost strategy produced strong T and B responses to HBV in various preclinical models of CHB (33–35). A combination of an HBsAg/HBcAg protein prime with TLR9 agonists followed by a boost with non-replicating modified vaccinia Ankara viruses, optimized for higher level expression of the same antigens was able to overcome the high-level tolerance observed in HBV-transgenic mice (35).

Given the evidence of high efficacy in at least 40% of the chronically infected mice, we plan to develop the VLV-based immunotherapy further and to initiate clinical trials to demonstrate safety and efficacy in CHB patients. Further improvement of VLV-based immunotherapy may include establishing a prime-boost strategy and/or combination with nucleos(t)ide treatment to lower HBV replication.

## Materials and Methods

### Plasmids and cells for VLV production

Plasmids for generation of empty vector VLV and the VLV expressing MHBs (MT2A) have been previously described (13, 14). We custom synthesized DNA fragments encoding HBV Pol and HBcAg with 2A peptides and cloned them into the MT2A plasmid using conventional cloning and/or Gibson assembly at SbfI and PacI restriction sites. We used BHK-21 cells for generation of VLV master stocks by Lipofectamine-mediated transfection and production of VLV working stocks by infection. We maintained BHK-21 cells in Dulbecco’s modified Eagle’s medium (DMEM) supplemented with 5% fetal bovine serum (FBS), 50 U/ml of penicillin, and 2 mM L-glutamine and switched them to Opti-MEM I reduced serum medium for VLV production for 48 to 72 h after infection. After preclearing of the conditioned medium by centrifugation at 600g for 10 min, we concentrated working stocks by ultrafiltration using MacroSep^®^ Advance 100 MWCO (Pall Laboratory) or Amicon Ultra 100K MWCO (EMD Millipore) centrifugal filter units. To quantify VLV titers, we used plaque assay or indirect immunofluorescence for VSV-G in BHK-21 as previously described (13, 14).

### Indirect immunofluorescence microscopy and flow cytometry

We seeded BHK-21 cells on coverslips one day prior to infection with VLVs at MOI=1 plaque forming unit (PFU)/cell. We fixed cells with 3% paraformaldehyde at 20 h post infection, washed with PBS containing 10 mM glycine (PBS-glycine), permeabilized with 0.1% Triton X-100 in PBS-glycine, and incubated with a 1:200 dilution of mouse monoclonal antibodies for MHBS (preS2, clone S 26), HBV Pol (clone 2C8) from Santa Cruz Biotechnology, VSV-G (clones I1 and I1-4), or rabbit polyclonal for HBcAg (DAKO). After extensive washes, we stained the cells with 1:500 dilution of goat anti-mouse AlexaFluor^®^ 488 IgG or goat anti-rabbit AlexaFluor^®^ 594 (ThermoFisher). We mounted washed coverslips on slides using ProLong Gold antifade reagent with DAPI (ThemorFisher) and imaged with Leica DMIRB microscope with 20×/0.75 objective, DC350FX camera (Leica) and ImagePro software or Nikon Eclipse 80i epifluorescence microscope with Retiga 2000R camera and NIS Elements software.

To validate expression of antigens by flow cytometry, we seeded BHK-21 cells in 12-well or 6-well tissue culture plates one day prior to infection with VLVs at MOI=1 plaque forming unit (PFU)/cell. We harvested cells at 20 h post infection with TrypLE Express (ThermoFisher), washed in FACS buffer (3% FBS in PBS supplemented with 0.1% sodium azide), treated with fixation and permeabilization buffer (ThermoFisher), and incubated with the PreS2, Pol, HBcAg and VSV-G antibodies listed above followed by rat monoclonal anti-mouse IgG1 conjugated to PE-Cy7 (BioLegend), goat anti-mouse IgG2a conjugated to AlexaFluor^®^ 647 and goat anti-rabbit conjugated to AlexaFluor^®^ 488 (ThermoFisher) antibodies. We acquired a minimum of 10,000 cells satisfying live cell forward and side scatter parameters using LSR II flow cytometer at the Cell Sorter Core Facility (Yale University School of Medicine) and analyzed the data with FlowJo software (version 7.6.5).

### Immunoblotting

We seeded BHK-21 cells in 100-mm tissue culture plates one day prior to infection with VLVs at MOI=1 plaque forming unit (PFU)/cell. We prepared protein lysates by scraping the cells in cold lysis buffer containing 1% Igepal (Sigma) and Complete^TM^ protease inhibitor cocktail and preclearing by centrifugation at 14,000 g for 10 min at 4°C. We separated proteins by SDS-PAGE in 4%–15% precast gradient gels (Biorad), transferred them onto nitrocellulose membranes, blocked, and incubated with rabbit polyclonal anti-2A-peptide (EMD Millipore), anti-VSV-G or anti-actin (SantaCruz) antibodies and HRP-conjugated or near infrared fluorescently labeled secondary antibodies. We scanned and analyzed immunoblots using ChemiDoc (Bio-Rad) or Odyssey imaging system (LICOR).

### In vivo studies (immunogenicity studies and models of HBV challenge and chronic infection)

All studies with animals were designed and carried out according to the recommendations in the Guide for the Care and Use of Laboratory Animals of the U. S. National Institutes of Health, Eighth Edition. IACUCs of University of Connecticut Health Center, Yale University, Albany Medical Center and Southern Research approved the studies carried out at the corresponding institutions. For immunogenicity studies, we used 6-8 week-old C57BL/6J male mice (Jackson Laboratory) at Yale University and CaroGen laboratory at UCHC TIP facility. We immunized animals with 10^8^ PFU/mouse in 1 ml of PBS via i.p. route and collected spleens from euthanized mice at day 7 post immunization. Control mice received equal volume of PBS at the time of immunization. We prepared single cell suspension of splenocytes by ACK lysis buffer, counted cells and stained with directly conjugated antibodies for T cell markers (CD4-Pacific Blue, CD8-AlexaFluor^®^ 488, CD62L-PE, CD44-AlexaFluor^®^ 647) in the presence of FcBlock (BD Biosciences). We also stimulated 10^6^ splenocytes per well with custom synthesized HBV peptides (Genscript) in the presence of 1X Brefeldin A (BioLegend) for 6 to 12 h prior to fixable viability staining eFluor-780 and surface staining with CD4-Pacific Blue, CD8-AlexaFluor^®^ 488. After extensive washes and treatment with fixation and permeabilization buffer (ThermoFisher), we stained cells with PE-conjugated anti-mouse IFNγ and AlexaFluor^®^ 647-conjugated anti-mouse TNF. We acquired a minimum of 50,000 cells satisfying live cell forward and side scatter parameters using LSR II flow cytometer at the Cell Sorter Core Facility (Yale University School of Medicine) and analyzed the data with FlowJo software (version 7.6.5). The H-2^b^ peptides used for CD8^+^ T cell stimulation were as follows: HBP-44 (NLNVSIPWTHKVGNF), HBP-396 (FAVPNLQSL), HBP-419 (VSAAFYHLPL), HBC-93 (MGLKFRQL), HBS-353 (VWLSVIWM), HBS-371 (ILSPFLPL). As a control peptide, we used H-2^d^ restricted peptide from HBV Pol (HYFQTRHYL). The threshold for cytokine positive CD8^+^ T cells was manually set using similarly gated and stimulated CD8^+^ T cells from spleens of non-immunized mice.

For acute challenge with HBV-AAV, we used 6-8 week-old CB6F1/J male mice (Jackson Laboratory) at Albany Medical College. We chose the CB6F1/J strain (H2^d/b^ haplotype) over C57BL/6 strain (H2^b-^ haplotype) for the HBV challenge because they generate a response to a dominant protective epitope of the surface antigen (HBS-191) restricted to H2^d^ haplotype that facilitates HBV clearance. We immunized animals with 10^8^ PFU/mouse via the i.p. route six weeks prior to delivery of 1011 genome copies of the AAV2/8 vector carrying 1.3 copies of the HBV genome (genotype D) via the i.v. route. Frequency of IFNγ-producing cells was measured by ELISPOT assay as previously described (36).

We established experimental chronic HBV replication by transduction of 6-8 week old C57BL/6J male mice in the facility of Southern Research (Birmingham, AL) following description of the model (27). After the HBV infection was confirmed by rising levels of serum HBsAg, we formed experimental groups with 10 animals per group based on normalization of HBsAg levels at day 21 post infection such that each group had the same mean and standard-deviation of 70±29 ng/mL HBsAg. Therefore, each group had a very similar range of HBsAg levels just prior to administration of VLVs. We immunized mice with 3xT2A or empty vector VLV (108 PFU/mouse) via i.p. route at day 28 post infection. We continued monitoring HBsAg levels for additional 13 weeks. On days 65 and 84, we administered plasmid DNA adjuvanted with 20 μg/mouse monophosphoryl lipid A from *Salmonella minnesota* (Invivogen) via the i.m. route as follows: the group of mice immunized with 3XT2A received three separate plasmids expressing HBsAg, HBV Pol and HBcAg in pVAC1 vector (Invivogen, 50 μg/mouse ea.); the group of mice immunized with empty vector VLV received empty pVAC1 vector (150 μg/mouse). We cloned HBsAg, HBcAg and Pol into the pVAC1 vector (Invivogen) and used EndoFree plasmid maxi kit (QIAGEN) to isolate DNA for the booster immunizations. To measure liver HBV RNA, we euthanized 5 mice per group to collect liver samples on day 70 and at the conclusion of the study on day 119. The numbers of IFNγ-producing CD8^+^ T cells were determined similarly to the analyses of immunogenicity in the naïve mice, except that HBV Pol and HBsAg antigen peptides were pooled for each antigen separately and no peptide control was used for background determination.

### Analytical assays for serum HBV antigens and liver HBV RNA

We collected blood samples from anesthetized mice weekly to measure serum HBsAg and HBeAg in the acute challenge experiment using ELISA kits from International Immunodiagnostics as previously described (14). We used HBsAg ELISA kit from XpressBio and the HBsAg standard from CellBioLabs to measure serum HBsAg in the chronic HBV infection model.

We collected liver samples from euthanized mice, snap-froze them in liquid nitrogen, and stored at −80° C or on dry ice until processing for RNA isolation. We isolated total RNA using the RNeasy Mini kit (QIAGEN) and used High Capacity cDNA Reverse Transcription kit (Applied Biosystems) for cDNA synthesis. We used quantitative TaqMan^TM^ Fast Advanced Master Mix (Applied Biosystems) and TaqMan^TM^ Assay Mix containing previously described probe and specific primers for detection of HBV: probe, 6FAM-5’ CCT CTT CAT CCT GCT GCT ATG CCT CAT C 3’-MGBNFQ, forward primer 5’-GTG TCT GCG GCG TTT TAT CA-3’, and reverse primer 5’-GAC AAA CGG GCA ACA TAC CTT-3’, and mouse GAPDH as an endogenous loading control on a StepOnePlus real time PCR system (Applied Biosystems) using StepOne software v2.3 (37). We used the comparative ΔCT method to determine HBV RNA abundance relative to mouse GAPDH mRNA.

### Statistical analyses

To determine the difference between the experimental groups, we used 1-way ANOVA for endpoint analyses (HBV RNA) or 2-way ANOVA for analyses over the time course (HBsAg) with Sidak’s multiple comparison test. To perform all calculations we used GraphPad Prism software, version 7 (GraphPad Software, San Diego, CA).

## Acknowledgements

We thank Lorraine Apuzzo, Mehdi Pouresmail, and Seyedehfatemeh Fazlhashemi at CaroGen Corporation for assistance with DNA cloning, production of VLV and characterization of the VLVs. We are also grateful to Drs. Raj Kalkeri, Kevin Walters, Krista Salley, Elizabeth Janci Wonderlich, and Jennifer Carl at Southern Research for their efforts on the chronic AAV-HBV model and to Shuguang Huang at Stat4ward for assistance with statistical analyses. This study was funded by the Connecticut Innovations, Connecticut Bioscience Innovation Fund, Gerpang Healthcare, and NIH grants R43DK113858 (V.N.) and R01AI124006 (M.D.R.). The content is solely the responsibility of the authors and does not necessarily represent the official views of the National Institutes of Health. The funders had no role in study design, data collection and interpretation, or the decision to submit the work for publication.

